# The bacterial microbiome in Spider and Deathwatch beetles

**DOI:** 10.1101/2024.07.15.603335

**Authors:** Austin Hendricks, T. Keith Philips, Tobias Engl, Rüdiger (Rudy) Plarre, Vincent G. Martinson

## Abstract

The beetle family Ptinidae contains a number of economically important pests, such as the Cigarette beetle *Lasioderma serricorne*, the Drugstore beetle *Stegobium paniceum*, and the diverse Spider beetles. Many of these species are stored product pests which target a diverse range of food sources from dried tobacco to books made with organic materials. Despite the threat that the 2,200 species of Ptinidae beetles pose, fewer than 50 have been surveyed for microbial symbionts, and only a handful have been screened using contemporary genomic methods. In this study, we screen 116 individual specimens that cover most subfamilies of Ptinidae, with outgroup beetles from closely related families Dermestidae, Endecatomidae, and Bostrichidae. We used 16S ribosomal RNA gene amplicon data to characterize the bacterial microbiomes of these specimens. The majority of these species had never been screened for microbes. We found that, unlike in their sister family Bostrichidae that has two mutualistic bacteria seen in most species, there are no consistent bacterial members of ptinid microbiomes. For specimens which had *Wolbachia* infections, we did additional screening using multilocus sequence typing, and showed that our populations have different strains of *Wolbachia* than has been noted in previous publications.

**Importance:** Ptinid beetles are both household pests of pantry goods and economic pests of dried good warehouses and cultural archives such as libraries and museums. Currently, the most common pest control measures for ptinid beetles are phosphine and/or heat treatments. Many ptinid beetles have been observed to have increasing resistance to phosphine, and heat treatments are not appropriate for many of the goods commonly infested by ptinids. Pest control techniques focused on symbiotic bacteria have been shown to significantly decrease populations, and often have the beneficial side effect of being more specific than other pest control techniques. This survey provides foundational information about the bacteria associated with diverse ptinid species, which may be used for future control efforts.

## Introduction

Many insects have expanded their niche space by developing close, codependent mutualisms with microbes (Cornwallis et al. 2023). These relationships are generally split into two categories: facultative and obligate. While facultative symbionts provide diverse benefits of protection and nutrition, nearly all obligate symbionts provide nutritional benefits – but identifying the benefits provided by individual symbionts requires experimental manipulation. An alternative method to help predict their functional category can be observed by screening individuals –symbiont types generally differ in their prevalence within and between host populations. Facultative symbionts can be either acquired from the parent (i.e., vertical inheritance) or from the environment (i.e., horizontal inheritance). While facultative mutualists provide benefits to the host under certain conditions (e.g., parasitoid pressure, heat stress), the cost to maintain the symbiont may leave hosts at a disadvantage when they no longer encounter the condition (Burke et al. 2010, Oliver et al. 2014). Therefore, facultative symbioses are not found in all populations of a host species or even all individuals within a population (Perreau and Moran 2022). In contrast, obligate mutualists are found in all host individuals because both host and symbiont rely on the other for survival, and closely related hosts generally have closely related symbionts as a result of co-diversification. These characteristics differentiate obligate and facultative symbionts and can be used in surveys of novel taxa to identify potential symbiotic relationships.

Obligately mutualistic bacterial symbionts have evolved independently at least 16 times and are found in 7 orders of insects: Blattodea, Coleoptera, Diptera, Hemiptera, Hymenoptera, Phthiraptera, and Psocoptera (Cornwallis et al. 2023); yet even more may be waiting to be identified. Beetles (Coleoptera) represent one of the most speciose group of insects, with over 350,000 described species, rivaled only by Hymenoptera (Farrell 1998, Bouchard et al. 2017, Forbes et al. 2018). Several beetle lineages are known to harbor microbial symbionts that include intracellular and extracellular bacteria and fungi (Salem & Kaltenpoth 2022).

One superfamily within Coleoptera that is responsible for a large number of timber and stored product pests is Bostrichoidea, which includes the Powderpost beetles (Bostrichidae), the Skin beetles (Dermestidae), and the Spider and Deathwatch beetles (Ptinidae). These lineages are specialists on difficult to digest, low-nutrient, low-moisture substrates. While many survive on dead wood, others have become specialists in a wide range of human products, ranging from processed foods like flour or herbs to culturally important artifacts like books, textiles, wool carpets, clothing, and furniture, making them an issue in archives and libraries (Hartnack 1939).

Bostrichoidea pests include: *Rhyzopertha dominica* (Lesser grain borer), that is primarily a pest of processed grain (Edde 2012) but has also been observed to directly affect agricultural yields through infestation of olive trees (Buonocore et al. 2017); *Trogoderma granarium* (Khapra beetle) the world’s most destructive pest of stored grains and seeds which is a quarantine species in Europe (Athanassiou et al. 2019), and *Prostephanus truncatus* (Larger grain borer), a major pest of maize that affects up to thirty percent of crops in central American and Africa (Quellhorst et al. 2021). Furthermore, Bostrichidae are notable because at least twenty-eight species were recently found to harbor two bacterial endosymbionts: *Candidatus* Bostridichola and *Candidatus* Shikimatogenerans (Okude et al. 2017, Engl et al. 2018, Kiefer et al. 2023). These endosymbionts encode pathways for the synthesis of nutrients essential for cuticle development and hardening, which allows these stored product pests to persist in the extremely dry environments of product warehouses (Engl et al. 2018, Kiefer et al. 2023). The remaining families of Bostrichoidea also contain specialized pests: Skin beetles (Dermestidae) that can survive in carpets and are used to strip skeletons of tissue, Spider beetles (Ptinidae s.s.) that common pests of stored products and dried goods, and Deathwatch beetles (Anobiidae s.s.) that are mainly associated with wood but contains major dried product pests. Many of these species demonstrate similar resistance to desiccation as Powderpost beetles (Yoder et al. 2009); yet despite similarities in natural history and economic importance as pests, there have been no systematic studies for obligate endosymbionts that may aide them in desiccation resistance or nutrient supplementation.

Ptinidae is comprised of 230 genera and 2200 species; however, there are likely many undocumented species because these beetles are difficult to collect due to their small size and that their larvae are hidden inside dietary compounds (e.g., wood) (Bell and Philips 2012). Broadly, the lineage can be split into two groups: the Spider beetles (subfamilies Ptininae and Gibbiinae), and the Deathwatch beetles (formerly Anobiidae; now the remaining 9 subfamilies of Ptinidae) (Lawrence 1991, Lawrence and Viedma 1991). Although some Deathwatch beetles were examined for endosymbionts by microscopy in the 1960s (Buchner 1965), no Spider or Deathwatch beetles have been confirmed to have bacterial endosymbionts. Furthermore, only five species of Deathwatch beetles have been confirmed to have *fungal* nutritional endosymbionts: *Lasioderma serricorne* (Cigarette beetle), *Stegobium paniceum* (Drugstore beetle), *Xestobium plumbeum*, and at least two species in the genus *Ernobius* (Jurzitza 1979). Strikingly, these fungal endosymbionts are from distantly related subphyla (Saccharomycotina & Pezizomycotina - most recent common ancestor >500 Mya), indicating that fungal symbionts have evolved independently multiple times (Martinson 2020). Moreover, these fungal symbionts may have been replacements for an ancestral bacterial symbiont related to that found in common ancestor of all Bostrichoidea.

Here we screen 116 individual specimens representing 61 species of Powderpost, Skin, Spider, and Deathwatch beetles for possible bacterial endosymbionts using high-throughput 16S ribosomal RNA gene high-throughput amplicon sequencing. We set out to answer (i) if there is an ancestral bacterial endosymbiont and (ii) describe the taxonomic composition of their bacterial microbiomes.

## Materials and Methods

### Sampling

Specimens came from a variety of sources. For 71 specimens, we used previously extracted DNA (Bell and Philips 2012, Gearner 2019). These specimens were collected from a wide geographic range across four different continents. For individual sample information, see Supplementary Table 1. The remaining 55 samples were sourced from insect laboratory colonies. The Spider beetles (*Gibbium aequinoctiale, Mezium affine, Mezium gracilicorne, Sphaericus gibbosus*) were sourced from the lab of TK Philips. The Deathwatch beetle *Anobium punctatum* and the Spider beetles *Gibbium psylloides* and *Niptus hololeucus* were sourced from the labs of T Engl and R Plarre. The Deathwatch beetles *Lasioderma serricorne* and *Stegobium paniceum* each had one population sourced from the USDA (E Scully, ARS, Plains Area, Manhattan, KS). The Skin beetle, *Dermestes maculatus*, and additional *S. paniceum* and *L. serricorne* were sourced from colonies maintained at the University of New Mexico (Albuquerque, NM). Full sample information is available in Supplemental Table 1.

Whole beetles were processed by extracting DNA from individuals. Specimens were either preserved in 100% ethanol or freeze dried. Ethanol-preserved specimens were dried for 1 hour prior to processing in a sterile biosafety cabinet. Individual specimens were processed with the Qiagen DNeasy Blood & Tissue Kit (Qiagen, Hilden, Germany). Briefly specimens were placed in RLT buffer and homogenized with a micro pestle followed by further homogenization using a Qiagen TissueLyser for 3 minutes at 30 Hz with 100 μL of 0.1 mm Mini-BeadBeater Zirconia-Silicate beads (BioSpec Products, Oklahoma, USA). The remaining steps were done following the manufacturer’s protocol. Library preparation and sequencing was done at Novogene (Beijing, China) using primers 515F (GTGCCAGCMGCCGCGGTAA) and 806R (GGACTACHVGGGTWTCTAAT), which target the V4 region of the 16S ribosomal RNA gene. Of the original set of 126 samples, 116 passed library preparation, representing 61 species. Sequencing was done on a 250 bp paired-end NovaSeq 6000 flowcell.

### Data QC & 16S rRNA Analysis

Read counts ranged between 30,359 – 118,987 with a mean of 85073 and a median of 90263. Denoising was done through dada2 with a forward truncation length of 225 bp and a reverse truncation length of 224 bp. Representative sequences were generated by Qiime2 (Bolyen et al. 2019), and taxonomic designations were made by aligning representative sequences to the Silva 138 database (Quast et al. 2013, Yilmaz et al. 2014). Representative sequences that had top alignments to either chloroplasts or eukaryotic mitochondria were removed from the dataset. The full count table and representative sequences file can be found in Supplemental Data 1 and Supplemental Data 2, respectively. Bar plots were generated using the qiime2R package (Bisanz 2018). Species richness and diversity indices were calculated using the *vegan* R package (Okasanen et al. 2022). NMDS plots were also created using the vegan and then rarefied using the *rrarefy* function to 16000 reads (Okasanen et al. 2022).

### Wolbachia Typing

To characterize the *Wolbachia* strains identified by high-throughput sequencing, we performed additional PCR and sequencing on *L. serricorne* and *S. paniceum* samples focusing on five housekeeping genes commonly used for *Wolbachia* multilocus sequence typing (MLST): *hcpA, gatB, ftsZ, coxA*, and *fbpA* (Baldo et al. 2006). Amplification was done with primers and cycling conditions utilized in previous studies (Baldo et al. 2006, Li et al. 2015). Amplicons were sent for Sanger sequencing at Eurofins (Luxembourg City, Luxembourg) or for linear amplicon sequencing by Oxford Nanopore at Plasmidsaurus (Oregon, USA). Sequences were then entered into the *Wolbachia* MLST database where they were assigned an allele number.

## Results and Discussion

### Overall microbiome diversity

Of the 116 specimens successfully sequenced, most were Spider and Deathwatch beetles, and the final dataset also included 5 Powderpost and 4 Skin beetles. We began by calculating overall diversity metrics for each family and subfamily of beetles. There was considerable variation in all diversity metrics amongst individuals. When grouped by subfamilies, Powderpost beetles had the highest alpha diversity of all groups (Figure 1a). There were no obvious trends with the average Shannon diversity indices and evenness for each subfamily (Figure 1b, 1c) although some individuals like the single endecatomid and single dryophilinid had high metrics for each. Anobiinae, Ptininae, and Xyletininae all include beetles reared in colonies as well as wild-caught beetles. We hypothesized that some of the variation seen in these subfamilies could be explained by these environmental variables, and re-ran the diversity statistics with these two groups differentiated (Figure 2a-c). As expected, wild-caught beetles had higher values for all three diversity indices used.

**Figure 1:**
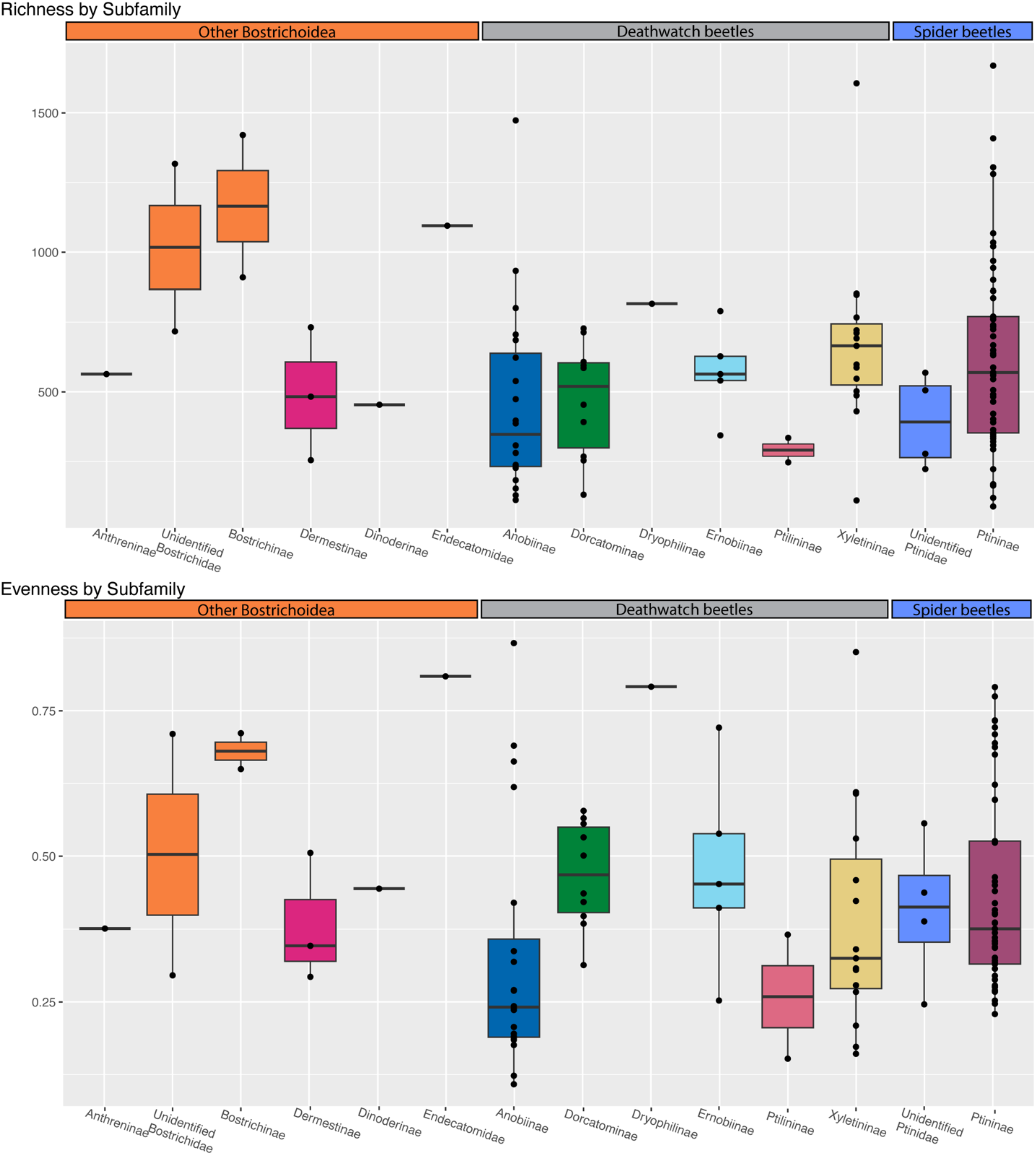
Barplots of diversity metrics by subfamily (or family when subfamily was not identified).B A) Richness/alpha diversity, B) Evenness, C) Shannon Diversity Index.

**Figure 2:**
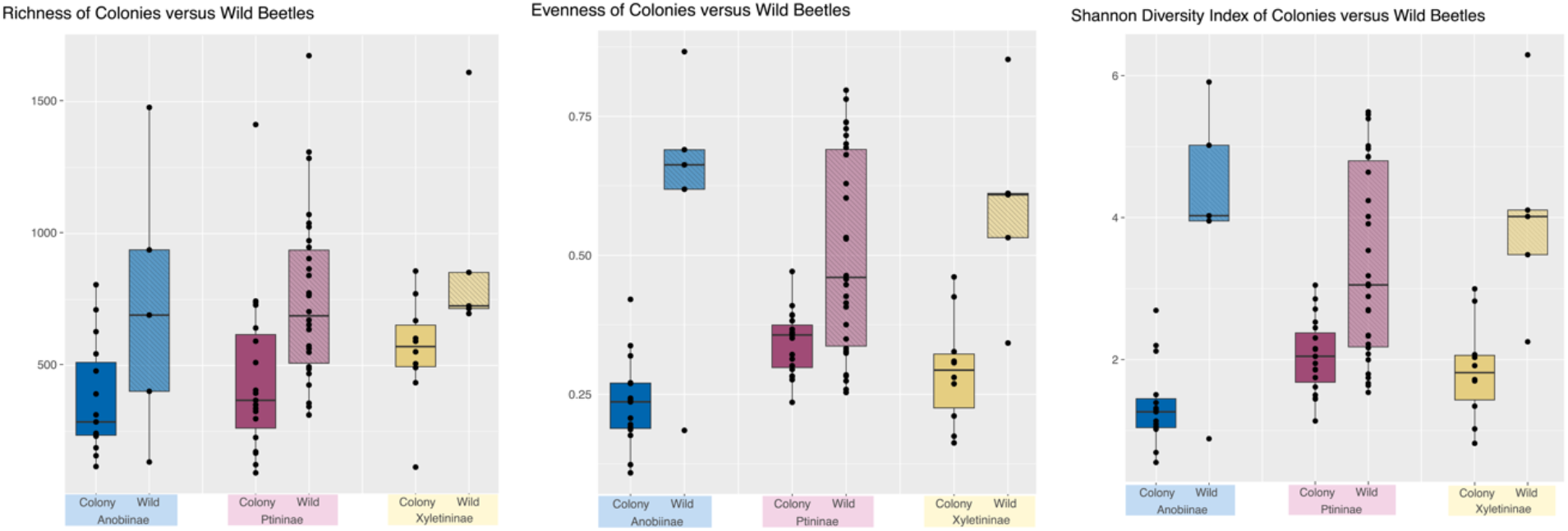
Barplots of diversity metrics for three subfamilies with specimens that are either wild caught or from a colony. A) Richness/alpha diversity, B) Evenness, C) Shannon Diversity Index.

We then compared overall microbiome composition using NMDS plots (Figure 3). There was significant overlap between all four families included in the dataset, but bostrichids and dermestids formed clusters in composition space (Figure 3a, stress = 0.0099). The endecatomid clustered near the bostrichids (Figure 3a). Conversely, Ptinidae specimens seemed to fall into two distinct clusters. To address these clusters, we tried grouping the points by subfamily instead of family (Figure 3b). In the left cluster are members of Dorcatominae and Ptilininae. In the right cluster is almost all Xyletininae. However, members of Ptininae and Anobiinae can be found in both clusters, albeit with different species on each side. This suggests that while some individuals within a species or within a subfamily do have similarities in their bacterial communities, that these commonalities are perhaps the result of similar environments, and do not indicate a strong taxonomic tendency towards hosting any particular communities of symbionts. Much of the variation seen in ptinid beetles can also be explained by the large number of ptinid specimens relative to the other subfamilies. There are only 4 Skin beetles, for instance, and three of those were sourced from the same colony.

**Figure 3:**
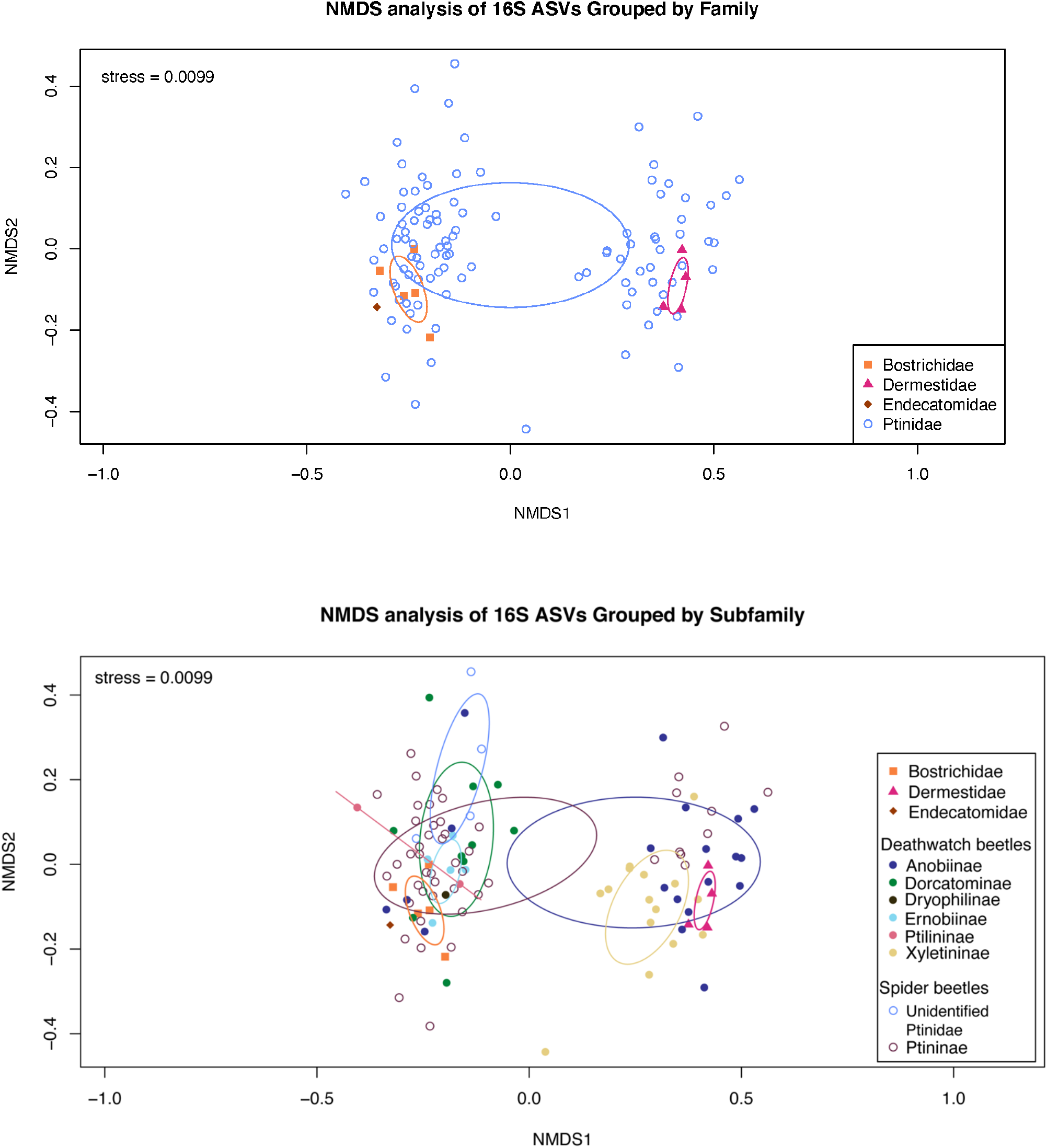
A) NMDS plot comparing similarities between individuals in each bostrichid family. B) NMDS plot comparing similarities between individuals based on families for outgroups (Bostrichidae, Dermestidae, and Endecatomidae) and subfamilies for ptinids. In cases where subfamily identification was not possible, the family is listed instead.

**Figure 4:**
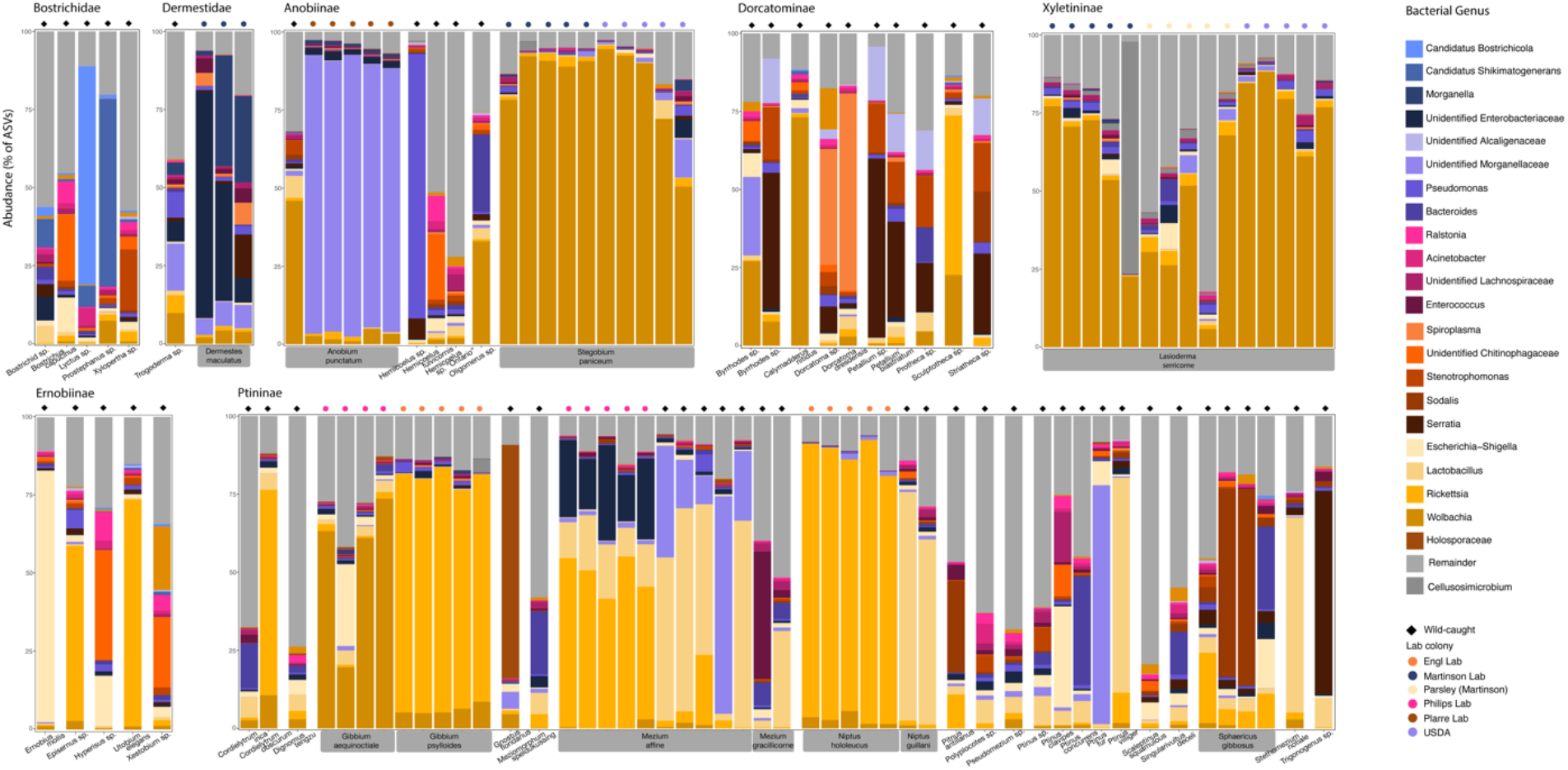
A stacked bar plot featuring the relative abundance of bacterial genera associated each individual beetle. Any bacterial genera that had less than 3% of the relative abundance for at least one individual were grouped together under “Remainder” for this plot. Bostrichids and dermestids are grouped by family, the remaining beetles are ptinids which are grouped by subfamily, and then spaced by genera. Source of the beetle is indicated by a small symbol on top of each bar plot, where shape indicates wild versus lab colony, and color indicates specific population. Species with more than one individual in the dataset are labeled with gray boxes beneath the bar plot.

### Powderpost & Skin beetles

Previous studies have shown that bostrichid beetles have one or both of the bacterial endosymbionts *Candidatus* Bostridichola and *Candidatus* Shikimatogenerans (Engl et al. 2018, Kiefer et al. 2023). In this dataset, the *Lyctus sp*. had ASVs corresponding to both *Bostridichola* and *Shikimatogenerans*, as expected. *Prostephanus sp*. had a small number of *Bostridichola* sequences, but was mostly dominated by *Shikimatogenerans* – possibly indicating that *Prostephanus* is in the evolutionary process of symbiont loss (*Bostridichola)*, similar to what was found in *Rhyzopertha dominica* (Kiefer et al. 2023). Alternatively, this may simply indicate low yield from the initial DNA extraction or bias during the PCR. Similarly, the *Xylopertha sp*. and *Bostrichus capucinus* specimens did not have abundant ASV’s for either endosymbiont, when at least *Shikimatogenerans* was expected from previous surveys (Kiefer et al. 2023).

Neither *Bostridichola* nor *Shikimatogenerans* were present in dermestids. Instead, *D. maculatus* individuals were dominated by sequences of *Morganella* and an unidentified Enterobacteriaceae. The *Morganella* sequence most closely aligned to *M. morganii* and *M. psychrotolerans*, which are an opportunistic pathogen and a species associated with food poisoning, respectively (Liu et al. 2016, Wang et al. 2020). All three *D. maculatus* individuals were sourced from a colony used to skeletonize mammal and bird specimens at the Museum of Southwestern Biology (MSB) at the University of New Mexico. Specimens being processed for accessioning to the MSB are sourced from diverse geographic locations. Because this *D. maculatus* colony subsists primarily on decomposing flesh with various origins, it is likely that either or both of these bacteria were acquired from their diet.

### Spider beetles

*Ptininae –* Ptininae samples were sourced from a mix of laboratory populations and wild-caught specimens. Unsurprisingly, specimens from the same source had similar microbiomes. While *Wolbachia* was present in both *Gibbium* species, *Rickettsia* was more common in Ptininae, with *Cordielytrum, Mezium, Niptus*, and *Sphaericus* all having sequences. *Mezium affine* and *Niptus guiliani* both had a significant portion of ASVs identified as *Lactobacillus rennini*, with *M. affine* from a lab population having over 10% of ASVs corresponding to *La. rennini* and wild *M. affine* and wild *N. guiliani* having over 50% of ASV’s corresponding to *La. rennini*. Other species in Ptininae also had some *Lactobacillus*, but this was generally 5% of ASVs or less. Of the 616 *Lactobacillus* ASV’s identified in the dataset, 333 were found in *Ptininae*.

The two populations of *Mezium affine* each had unique microbiome compositions, likely because they were maintained as colonies at different places with different diets. The *Mezium affine* lab colony had >50% of sequences corresponding to *Rickettsia*, whereas the wild-caught samples were more likely to have *Lactobacillus* or an unidentified Morganellaceae. *Sodalis*, an genus with many insect endosymbionts found in taxonomically diverse hosts (Tláskal et al. 2021), was identified in *Pitnus antillus* and *Sphaericus gibbous. Gnostus floridanus* was dominated by *Holosporaceae*. Finally, an unidentified *Trigonogenius* had over 65% of its ASV counts corresponding to the same ASV identified as a *Serratia*. While we cannot identify the species, the facultative symbiont *Serratia symbiotica* has been noted in aphids, where it confers a number of advantages, including protection from heat stress and parasitism (Perreau et al. 2021). Further investigation of the bacterial communities of *Trigonogenius* may be warranted.

### Deathwatch beetles

*Anobiinae –* Many samples in *Anobiinae* were sourced from lab colonies, and we found that individuals within the same colony had very similar microbiomes. While the wild *Anobium punctatum* was dominated by *Wolbachia* ASVs, the specimens from lab colonies had over 90% of ASVs correspond to an unidentified Morganellaceae endosymbiont. When the sequence was entered into a BLAST search, it had a high percent identity to *Symbiodolus* (CP152425.1), a recently described arthropod symbiont (Wierz et al. 2024).

There was no consistency in the microbiomes for *Hemicoelus*, with only *Pseudomonas* and an unidentified Chitinophagaceae standing out. While *Pseudomonas* has been associated with bark beetle microbiomes, it has also been reported as an entomopathogen (Saati-Santamaría et al. 2021). Unfortunately, *Pseudomonas* species and strains cannot be differentiated based on the 16S rRNA sequences included in this study.

For the *Oligomerus sp*., a *Bacteroides* sequence was identified. However, this sequence does not appear to be related to *Candidatus* Bostridichola or *Candidatus* Shikimatogenerans, and instead matched most closely to *Phocaeicola vulgatus*, which is associated with the human gut microbiome (Wang et al. 2021). *Oligomerus sp*. and *S. paniceum* additionally both had a large number of ASVs matching to *Wolbachia*.

*Dorcatominae – Byrrhodes, Petallium, Protheca*, and *Striatheca* all had an ASV identified as a *Stenotrophomonas* that comprised 13-16% of all ASVs for those specimens. One *Byrrohodes* specimen also had a high association (>25% of ASVs) with a different insect-associated Morganellaceae ASV than was seen in *A. punctatum* although it had similar BLAST results (99.21% identity to CP152425.1), indicating they may be the same species. Both *Dorcatoma* specimens had over 25% of ASVs correspond to the same *Spiroplasma* ASV, which had a 100% identity on BLAST to numerous *Spiroplasma* arthropod endosymbionts, such as one for ticks (LC388762.1), *Drosophila melanogaster* (PP593989.1), and the pea aphid *Acyrthosiphon pisum* (KP710442.1). Either *Wolbachia* or *Rickettsia* were also found in *Byrrhodes, Calymadderus*, and *Sculptotheca. Striatheca* additionally had 16% of ASVs correspond to a *Sodalis*.

*Ernobiinae –* The dominant bacterial genera in Ernobiinae were *Escherichia-Shigella, Rickettsia*, and an unidentified Chitinophagaceae.

*Xyletininae –* For this group, the only species represented is *Lasioderma serricorne*, which was sourced from three populations: a laboratory population maintained at UNM, a population sourced from the USDA, and a population sourced from a commercially sold jar of dried parsley in Miami, Florida. All specimens were infected with *Wolbachia*.

### Wolbachia Typing for L. serricorne and S. paniceum

*Wolbachia* infections have been previously reported in both *L. serricorne* and *S. paniceum*, and studies have utilized either *wsp* or MLST primers to genotype these endosymbionts (Kageyama et al. 2010, Li et al. 2015). We utilized the MLST primer set to survey our *L. serricorne* and *S. paniceum* populations, all of which harbored *Wolbachia* according to the 16S amplicon survey. These results varied in that individual genes were not amplified, or had a mixed chromatogram in the Sanger results, which is why the Parsley population was unable to be typed. This indicated that there might be co-infections of multiple *Wolbachia* types in some populations. From the genes that are available, it appears that the *L. serricorne* populations (USDA, UNM) each have a unique *Wolbachia* strain, but the UNM population of *L. serricorne* may match the Canadian *L. serricorne* population from the Li et al. 2015 study. The same *Wolbachia* strain seemingly infects the USDA and SOBA *S. paniceum* populations, but it is unique from the strain identified in Li et al. 2015 in *S. paniceum* from that are also from the US. Overall, *Wolbachia* infection and MLST types vary among populations, which together, indicate independent acquisition of this common endosymbiont. It is unlikely that *Wolbachia* plays an important role in Deathwatch biology; however, future experiments are required to test how eliminating *Wolbachia* affects host biology.

## Conclusion

Deathwatch and Spider beetles have extremely varied bacterial microbiomes with few commonalities aside from the reproductive manipulators *Wolbachia* and *Rickettsia*. Although the exact function of these symbionts is currently unknown, they likely represent facultative symbionts that may be reproductive manipulators or provide a context-dependent benefit their hosts (e.g., protection from viruses). This study does not support the existence of a consistent, mutualistic, obligate bacterial endosymbiont in all Bostrichoidea beetles. However, amplicon sequencing is limited by the length of the sequences and quality of the DNA. It is possible that these obligate symbionts do exist, but we were unable to identify them, or unable to amplify a sequence for them. This may also be why two of the Powderpost beetles in this study did not have the expected *Bostridichola* or *Shikimatogenerans*. Furthermore, while the currently recognized and described symbionts in Powderpost beetles (bacteria) and in Deathwatch beetles (fungal) provision hosts with nutrients used in development, they provide different metabolites: tyrosine and B-vitamins, respectively (Engl et al. 2018, Martinson 2020, Kiefer et al. 2023). This may suggest that the selective forces that created these symbiotic relationships occurred in each lineage independently following their divergence.

Obligate symbionts have been proposed as targets for new, specific pesticides, as many of the hosts of these obligate symbionts rely on them for the nutrients needed to complete development (Gupta et al. 2023). However, other symbionts are also potential targets, such as *Wolbachia*, which has already been used to control the population of mosquitos by inducing cytoplasmic incompatibility between treated mosquitos and wild mosquitos (Zheng et al. 2019). In other cases, engineered *Wolbachia* strains have been used to partially vaccinate insects against diseases that they serve as a vector for, as attempted in mosquitos against dengue and planthoppers against a rice virus (Guo et al. 2023). These techniques could potentially be used in the future to reduce beetle populations of the common pests – *L. serricorne* (the Cigarette beetle) and *S. paniceum* (the Drugstore beetle) – which both seem to have widespread *Wolbachia* infections, especially as phosphine resistance is becoming more common (Sağlam et al. 2015). Better understanding the microbiomes across the diversity of insects is critical for future control efforts and the development of new technologies to combat persistent, agriculturally and economically important pests.

---

**Table 1.** Table of MLST gene classifications from each study population, compared to examples from the literature.

| Publication | Species | Population | gatB | coxA | hcpA | ftsZ | fbpA | wsp |
| --- | --- | --- | --- | --- | --- | --- | --- | --- |
| Kageyama et. al. 2010 | *Lasioderma serricorne* | Ls1 | N/A | N/A | N/A | N/A | N/A | Single: Type 3 (16) |
| Li et. al. 2015 | *Lasioderma serricorne* | Canada | 4 | 14 | 40 | N/A | 4 | N/A |
| Li et. al. 2015 | *Lasioderma serricorne* | Spain | 4 | 14 | N/A | N/A | N/A | N/A |
| Li et. al. 2015 | *Lasioderma serricorne* | Greece | 96 | N/A | N/A | 73 | N/A | N/A |
| This paper | *Lasioderma serricorne* | UNM | N/A | 14 | N/A | N/A | 4 | N/A |
| This paper | *Lasioderma serricorne* | USDA | 4 | 14 | 40 | 243 | 444 | N/A |
| This paper | *Lasioderma serricorne* | Parsley | N/A | N/A | N/A | N/A | N/A | N/A |
| Kageyama et. al. 2010 | *Stegobium paniceum* | Sp1 | N/A | N/A | N/A | N/A | N/A | Double: Type 13 (4) & Type 14 (9) |
| Li et. al. 2015 | *Stegobium paniceum* | USA | N/A | 64 | N/A | 63 | 75 | N/A |
| This paper | *Stegobium paniceum* | UNM | 103 | 64 | 40 | 243 | 444 | N/A |
| This paper | *Stegobium paniceum* | USDA | 103 | 64 | 40 | 243 | N/A | N/A |

## Notes

### Competing Interest Statement

The authors have declared no competing interest.

